# Application of Cas12j for *Streptomyces* editing and cluster activation

**DOI:** 10.1101/2021.10.28.465406

**Authors:** Lee Ling Tan, Elena Heng, Nadiah Zulkarnain, Chung Yan Leong, Veronica Ng, Lay Kien Yang, Deborah Chwee San Seow, Guangrong Peh, Yee Hwee Lim, Koduru Lokanand, Yoganathan Kanagasundaram, Siew Bee Ng, Fong Tian Wong

## Abstract

In recent years, CRISPR-Cas toolboxes for *Streptomyces* editing have rapidly accelerated natural product discovery and engineering. However, Cas efficiencies are also oftentimes strain dependent, subsequently a variety of Cas proteins would allow for flexibility and enable genetic manipulation within a wider range of *Streptomyces* strains. In this work, we have further expanded the Cas toolbox by presenting the first example of Cas12j mediated editing in *Streptomyces* sp. A34053. In our study, we have also observed significantly improved editing efficiencies with *Acidaminococcus sp*. Cas12j compared to Cas12a, *Francisella tularensis subsp. novicida* U112’s type V-A Cas (FnCpf1).

## Introduction

Actinomycetes are highly productive factories of bioactive natural products (NP) [1, 2]. The activation and production of these pharmaceutically important, secondary metabolites are typically triggered by environmental signals [3]. Unfortunately, under laboratory conditions, it is estimated that 80 % of this chemical diversity is silent [4-6]. To this end, various methods to activate these silent biosynthetic gene clusters were investigated and deployed [7]. In our laboratory and others, CRISPR-Cas mediated editing of actinomycetes, in particular *Streptomyces*, have accelerated NP discovery and engineering [8-10].

Previously, *Streptococcus pyogenes* Cas9 (SpCas9), *Staphylococcus aureus* Cas9 (SaCas9), *Streptococcus thermophilus* CRISPR 1 Cas9 (Sth1Cas9) and *Francisella tularensis subsp. novicida* U112’s type V-A Cas (FnCpf1) have been used for genome editing in *Streptomyces* [8]. Cpf1, also known as Cas12a, has been shown to be highly efficient (75 - 95%) for precise genome editing in the presence of a homology repair template in *Streptomyces* [11]. This is comparable to the more commonly used SpCas9. In 2020, Pausch et al. reported the discovery and usage of the hypercompact Cas12 sub family *Acidaminococcus sp*. Cas12j (AsCas12j) where the Cas protein has a single RuvC endonuclease domain for both processing of crRNA and cleavage [12]. Although it has similar functions and capabilities as Cas12a, Cas12j is significantly smaller at 757 amino acid long, as compared to SpCas9 (1368 aa) and FnCpf1 (1300 aa). The AsCas12j system was demonstrated to be active *in vitro* and in human and plant cells.

Out of the three AsCas12j examined, AsCas12j-2 was reported to be self-processing and have high levels of editing efficiencies. Subsequently, we selected AsCas12j-2 for editing in *Streptomyces* sp. A34053 from the A*STAR Natural organism library (NOL) [13]. In this study, we demonstrated that AsCas12j-2 is functional in *Streptomyces* and in these examples, has even outperformed that of FnCpf1. This is the first reporting for the successful application of Cas12j in genome manipulation of *Streptomyces*.

## Methods

### Conjugation and screening of strains

Spore preparations, conjugation protocol as well as the method for the screening of edits were adapted from Yeo et al, 2019 [8].

To obtain the spores of *Streptomyces* sp. A34053, the strain is first propagated in SV2 media (For 1 L, add 15 g glucose, 15 g glycerol, 15 g soya peptone (Biobasic), 1 g CaCO_3_ to deionized water, pH 7.0) and plated onto ISP Medium No. 4 (BD Biosciences). Spores were removed and resuspended in sterile TX buffer (50 mM Tris pH 7.4, 0.001% (v/v) Triton X). Intergeneric conjugation between *Streptomyces* sp. A34053 and DNA methylase deficient WM3780 *E. coli* donor cells were performed by transforming promoter knock-in constructs into WM3780.

Apramycin-sensitive clones were picked for screening by colony PCR. Successful PCR amplicons were screened for the presence of the promoter or promoters via restriction enzyme digest. Positive samples were then validated with Sanger sequencing.

### Fermentation and analysis

Wild type *Streptomyces* sp. A34053 and edited mutants with promoter knock-in were cultured in 5 mL SV2 media for 3 - 5 days. Saturated seed cultures were diluted into fresh fermentation media: SV2, ISP2 (For 1 L, add 10 g malt extract (Sigma-Aldrich), 4 g yeast extract (BD Biosciences), and 4 g glucose.), and CA08LB (For 1 L, 15 g glucose, 20 g cane molasses, 40 g soluble starch, 8 g CaCO_3_, and 25 g cottonseed flour (Sigma-Alrich), pH adjusted to pH 7.2.) in a 1:20 volume ratio and fermented with 200 rpm shaking at 30 °C in the dark. After 9 days, the cultures were pelleted then the separated biomass and supernatant were lyophilized. The dried samples were extracted by methanol then filter through filter paper (Whatman Grade 4) and filtrates were reconstituted for analysis.

The extracts were analysed on an Agilent 1290 Infinity LC System coupled to an Agilent 6540 accurate-mass quadrupole time-of-flight (QTOF) mass spectrometer. 5 µL of extract was injected onto a Waters Acquity UPLC BEH C18 column, 2.1 × 50 mm, 1.7 µm. Mobile phases were water (A) and acetonitrile (B), both with 0.1 % formic acid. The analysis was performed at flow rate of 0.5 mL/min, under gradient elution of 2 % B to 100 % B in 8 min. Both MS and MS/MS data were acquired in positive electrospray ionization (ESI) mode. The typical QTOF operating parameters were as follows: sheath gas nitrogen, 12 L/min at 325 °C; drying gas nitrogen flow, 12 L/min at 350 °C; nebulizer pressure, 50 psi; nozzle voltage, 1.5 kV; capillary voltage, 4 kV. Lock masses in positive ion mode: purine ion at *m/z* 121.0509 and HP-0921 ion at *m/z* 922.0098.

## Results

### 1. Design of 12j vector

To utilize AsCas12j-2, the backbone of an all-in-one pCRISPomyces-2 plasmid was used (Addgene #61737 [14]), where the Cas protein was replaced with codon optimized AsCas12j-2 from Twist Biosciences (USA). The crRNA sequence [12], “5-GTCGGAACGCTCAACGATTGCCCCTCACGAGGGGAC-3”, was inserted after the promoter transcribing the guide RNA. The general annotation of the elements in this plasmid is shown in Figure 1A.

**Figure 1.**
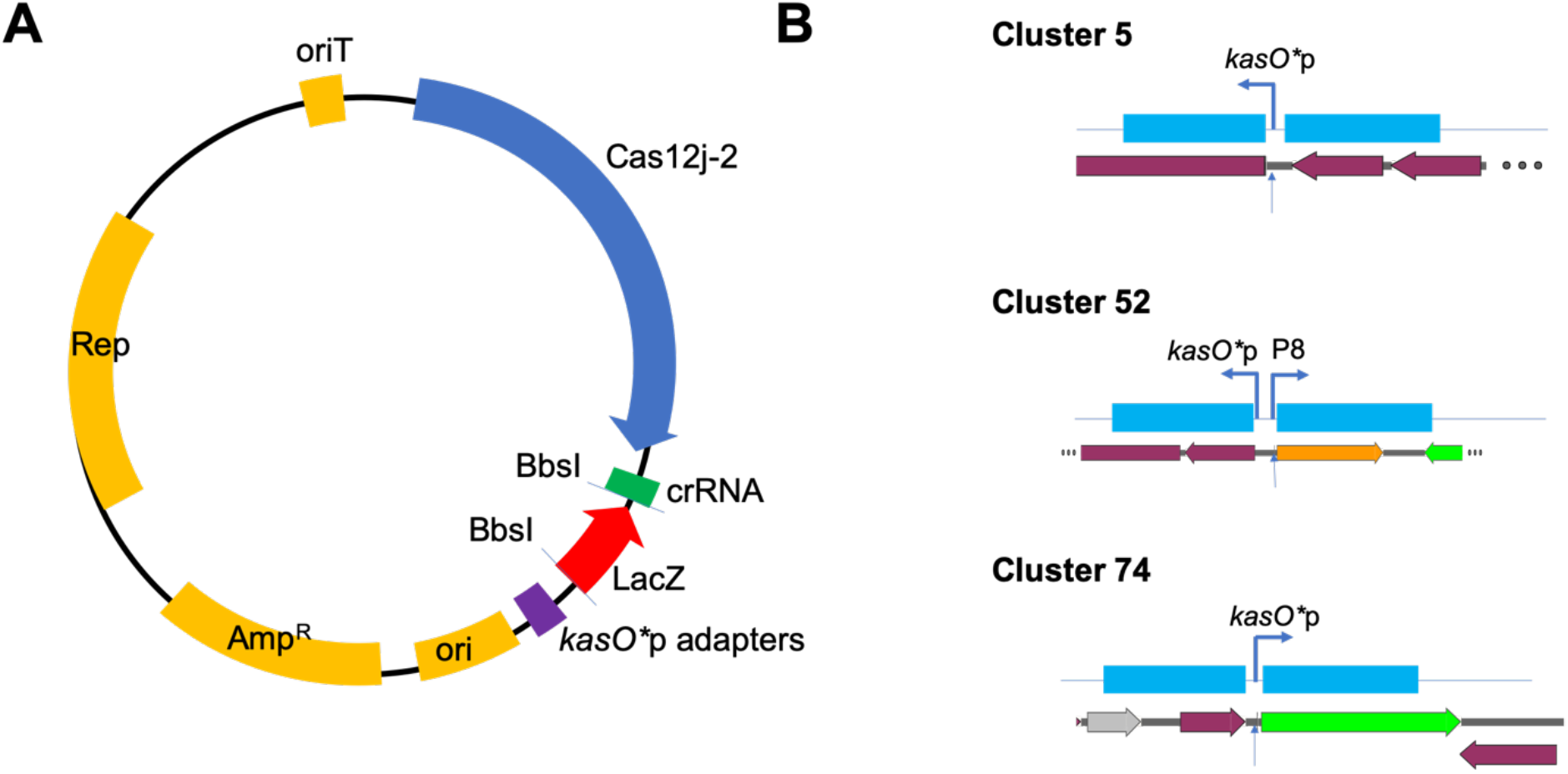
(A) General map of all-in-one editing CRISPR-Cas construct for one-step genome editing of *Streptomyces* using AsCas12j-2 along with (B) schematic homology flanks used for *Streptomyces* sp. A34053 editing. (A) Rep: Replicon, AmpR: aparamycin resistance cassette, ori: origin of replication, oriT: origin of transfer, LacZ: LacZ operon for screening, *kasO*\*p adapters for addition of homology flanks, crRNA: CRISPR-RNA, BbsI sites for golden gate assembly of protospacer. (B) Homology flanks (Blue) with insert of promoters are designed as the figure, where each flank is ∼2 kb in length and a 97 bps (*kasO*\*p) or 778 bps (*kasO*\*p with P8 promoter) is inserted at the specified sites. The sites of the protospacer are also indicated on the genome with arrows. Legend of genes on the genome annotation; Grey: domain of unknown function, orange: transporter, green: regulator, red: biosynthetic enzymes.

### 2. Functional Cas12j in comparison with Cas12a

Since FnCpf1 and AsCas12j-2 belong to the Type V CRISPR-Cas system and recognize similar PAM sites (TTN), and FnCpf1 was observed to be significantly successful in previous experiments [8], in this study, we will be comparing the genome editing capabilities of AsCas12j-2 with FnCpf1.

Three different clusters within *Streptomyces* sp. A34053 (Figure 1B) were targeted for promoter exchange to enable cluster activation [15]. Since FnCfp1 and AsCas12j-2 uses the same PAM site, the same protospacer (24 bps) and homology flanks were used for both Cas proteins. In the editing of all three clusters, AsCas12j-2 was functional and was able to insert the new promoter (Table 1). The edited strains also demonstrated changes in chemical profiles with the promoter exchange (Figure 2). This would be the first example of Cas12j in *Streptomyces*. Interestingly, both AsCas12j-2 and FnCpf1 gave similar numbers of exconjugants but AsCas12j-2 consistently has a significantly high rate of edits whilst FnCpf1 yield no edited strains (Table 1). This suggests that although AsCas12j-2 and FnCpf1 have similar toxicity within *Streptomyces* sp. A34053, AsCas12j-2 outperforms FnCpf1 in genome editing, within the context of these clusters.

**Figure 2.**
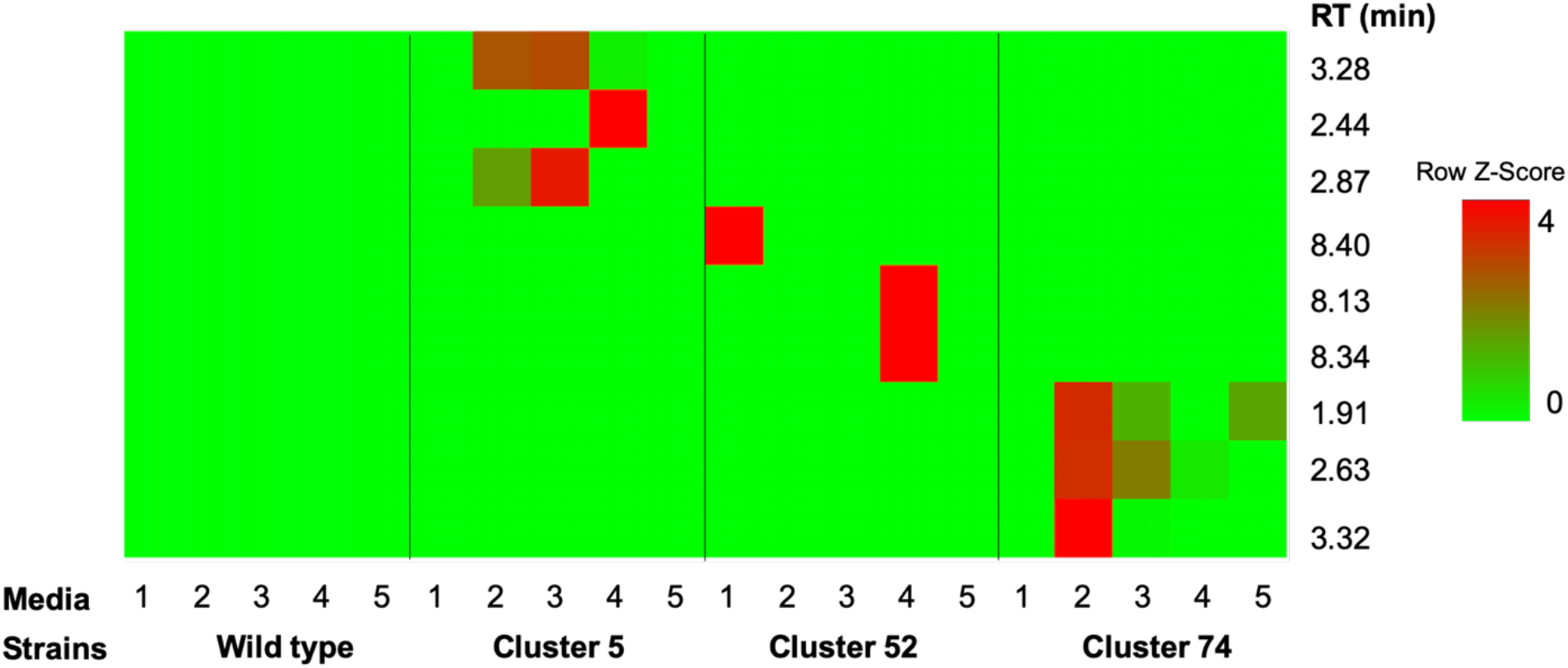
Heat map representation of secondary metabolites production between wild type and edited mutants for Cluster 5, 52 and 74 (Table 1). The comparison was done across 5 culture media (1: CA02LB, 2: CA07LB, 3: CA08LB, 4: CA09LB, 5: CA10LB). The heat map is generated by shinyheatmap interface [17] and scaled across rows.

**Table 1.** Editing efficiencies of AsCas12j-2 and FnCpf1.

| Cluster | Cas protein | Insertion (kb) | Efficiency <sup>a</sup> | Exconjugants <sup>b</sup> |
| --- | --- | --- | --- | --- |
| 5 | AsCas12j-2 | 0.1 | 50 % (3/6) | >50 |
|  | FnCpf1 | 0.1 | 0 % (0/8) | >50 |
| 74 | AsCas12j-2 | 0.1 | 75 % (3/4) | 4 |
|  | FnCpf1 | 0.1 | 0 % (0/6) | 6 |
| 52<sup>c</sup> | AsCas12j-2 | 1 | 75 % (3/4) | 7 |

<sup>a</sup> Editing efficiency given by the number of correct clones validated by sanger sequencing over the total number of clones picked (Supplementary Figure 1).
<sup>b</sup> Number of exconjugants observed per 100 µL of spore preparation used in each conjugation. A typical spore prep contains ~10<sup>6</sup> - 10<sup>7</sup> spores/mL as determined by serial dilution plating.
<sup>c</sup> Genome editing using FnCpf1 was not performed for this cluster.

## Discussion

To harness the full biosynthetic potential of *Streptomyces*, much efforts have been placed in expanding the Cas toolbox for activation of silent biosynthetic gene clusters. The Cas protein, its associated levels of toxicity, its PAM recognition sites, as well as its efficiency of editing the genome are all important factors of consideration in its application in *Streptomyces* [16]. Here we report the successful application of Cas12j for *Streptomyces* usage. AsCas12j-2 is not only functional in S*treptomyces* sp. A34053, but it also has a higher editing efficiency compared to FnCpf1 in this study.

We believe that the hypercompact AsCas12j-2 will be a valuable tool in our strategy to generating genomic perturbations in *Streptomyces*. Previous studies have also shown successful applications of various CRISPR-Cas strategies from *Streptomyces* to the rare Actinomycetes without modifications or addition of helper proteins. In this same manner, we predict that the usage of Cas12j-2 could also be applied across the Actinomycetes family.

## Acknowledgement

This work is supported by the National Research Foundation (NRF) 19^th^ Competitive Research Program (CRP) grant - NRF-CRP19-2017-05.

---

**Supplementary Figure 1.**
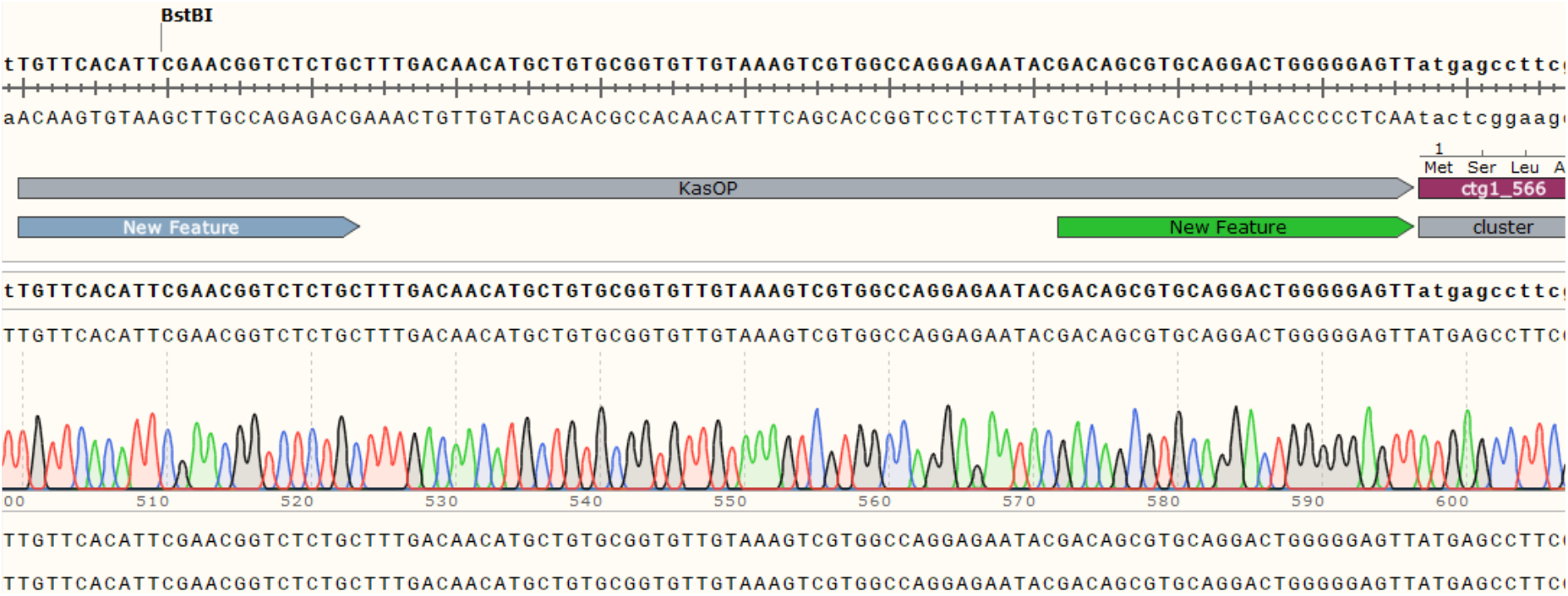
Representative trace of edited genome sequence (insertion of *kasO*\*p) for cluster 74 in S*treptomyces* sp. A34053.

